# RNA aptamer reveals nuclear TDP-43 pathology is an early aggregation event that coincides with *STMN-2* cryptic splicing and precedes clinical manifestation in ALS

**DOI:** 10.1101/2023.10.24.563701

**Authors:** Holly Spence, Fergal M. Waldron, Rebecca S. Saleeb, Anna-Leigh Brown, Olivia M. Rifai, Martina Gilodi, Fiona Read, Kristine Roberts, Gillian Milne, Debbie Wilkinson, Judi O’Shaughnessy, Annalisa Pastore, Pietro Fratta, Neil Shneider, Gian Gaetano Tartaglia, Elsa Zacco, Mathew H. Horrocks, Jenna M. Gregory

**Affiliations:** Institute of Medical Sciences, University of Aberdeen, Aberdeen, UK; EaStCHEM School of Chemistry, University of Edinburgh, Edinburgh, UK; IRR Chemistry Hub, Institute for Regeneration and Repair, University of Edinburgh, Edinburgh, UK; Department of Neuromuscular Diseases, UCL Queen Square Institute of Neurology, London, UK; Centre for Discovery Brain Sciences, University of Edinburgh, UK; RNA System Biology Lab, Instituto Italiano di Tecnologia, Genoa, Italy; NHS Grampian tissue biorepository, Department of Pathology, Aberdeen, UK; The Maurice Wohl Institute, King’s College London, London, UK; Department of Neurology, Center for Motor Neuron Biology and Disease, Columbia University, New York, USA

**Keywords:** TDP-43, Loss-of-function, Neuropathology, Cryptic splicing, *Stathmin-2*, RNA aptamer, Amyotrophic lateral sclerosis, Cognition

## Abstract

TDP-43 is an aggregation-prone protein which accumulates in the hallmark pathological inclusions of amyotrophic lateral sclerosis (ALS). However, analysis of deeply-phenotyped human *post-mortem* samples has shown that TDP-43 aggregation, revealed by standard antibody methods, correlates poorly with symptom manifestation. Recent identification of cryptic-splicing events, such as the detection of *Stathmin-2* (*STMN-2*) cryptic exons, are providing evidence implicating TDP-43 loss-of-function as a potential driving pathomechanism, but the temporal nature of TDP-43 loss and its relation to the disease process and clinical phenotype is not known. To address these outstanding questions, we used a novel RNA aptamer, TDP-43^APT^, to detect TDP-43 aggregation and used single molecule *in situ* hybridization to sensitively reveal TDP-43 loss-of-function and applied these in a deeply-phenotyped human *post-mortem* tissue cohort. We demonstrate that TDP-43^APT^ identifies pathological TDP-43, detecting aggregation events that cannot be detected by classical antibody stains. We show that nuclear TDP-43 pathology is an early event, occurring prior to cytoplasmic aggregation and is associated with loss-of-function measured by coincident *STMN-2* cryptic splicing pathology. Crucially, we show that these pathological features of TDP-43 loss-of-function precede the clinical inflection point and are not required for region specific clinical manifestation. Furthermore, we demonstrate that gain-of-function in the form of extensive cytoplasmic aggregation, but not loss-of-function, is the primary molecular correlate of clinical manifestation. Taken together, our findings demonstrate implications for early diagnostics as the presence of *STMN-2* cryptic exons and early TDP-43 aggregation events could be detected prior to symptom onset, holding promise for early intervention in ALS.

**Short Abstract:** Recent identification of cryptic-splicing events such as the detection of *Stathmin-2* (*STMN-2*) cryptic exons, are providing evidence implicating TDP-43 loss-of-function as a potential driving pathomechanism in amyotrophic lateral sclerosis (ALS). However, the temporal nature of TDP-43 loss and its relation to clinical phenotype is not known. Here, we used a novel RNA aptamer to detect TDP-43 aggregation and used single molecule ISH to sensitively reveal TDP-43 loss-of-function, applying these methods in a deeply-phenotyped human *post-mortem* tissue cohort. We show that nuclear TDP-43 pathology is an early event, that coincides with *STMN-2* cryptic splicing. Crucially, we show that these pathological features of TDP-43 loss-of-function precede the clinical inflection point and are not required for region specific clinical manifestation. Furthermore, we demonstrate that gain-of-function, but not loss-of-function, is the primary molecular correlate of clinical manifestation. Taken together, our findings demonstrate implications for early diagnostics and intervention prior to symptom onset in ALS.

**Graphical Abstract:** 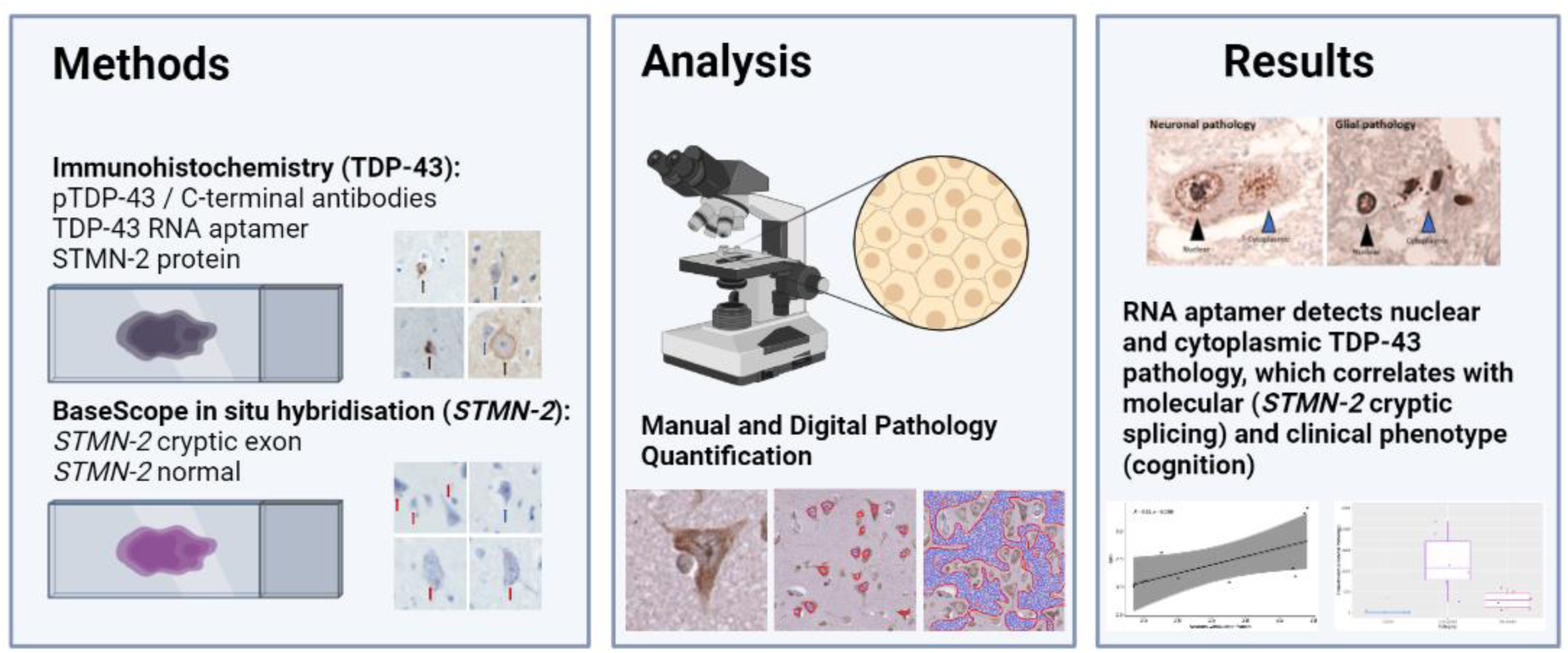

## Introduction

TDP-43, encoded by the *TARDBP* gene, is an RNA-binding protein that predominantly localizes to the nucleus, and whose mislocalization and accumulation in pathologically phosphorylated cytoplasmic aggregates is a hallmark of amyotrophic lateral sclerosis frontotemporal dementia spectrum disorders (ALSFTSD). Indeed, TDP-43 pathology is not just restricted to ALSFTSD, it can be seen in other neurodegenerative diseases such as limbic-predominant, age-related TDP-43 encephalopathy (LATE), and even in the brains of individuals without neurological deficits as a part of aging^1,2^. The hypothesized mechanisms underpinning the role of TDP-43 pathology in the pathogenesis of ALSFTSD can be broadly categorised by (i) gain-of-toxic function, owing to the presence of cytotoxic insoluble inclusions accumulating in the cytoplasm and (ii) loss-of-function as insoluble, TDP-43 can no longer carry out its normal cellular function.

TDP-43 is known to function as a repressor of cryptic exons during splicing^3,4^. TDP-43 represses cryptic exon inclusion in *Stathmin-2 (STMN-2)* through its binding site in the first intron, but when pathologically mislocalized, TDP-43 fails to repress the incorporation of cryptic exon 2A into mature mRNA^5^. Truncated *STMN-2* mRNA subsequently arises because of a premature polyadenylation signal in cryptic exon 2A itself^6–8^. Thus, detection of this incorrectly spliced *STMN-2* cryptic exon is a sensitive molecular marker of TDP-43 loss-of-function. Indeed, loss-of-function mechanisms have been demonstrated to play an important role in disease pathogenesis in preclinical^9–11^ and clinical studies^12^. Additionally, evidence exists supporting gain-of-function mechanisms underpinning the pathogenic role of TDP-43 in ALSFTSD. We have shown previously that the presence of phosphorylated TDP-43 (pTDP-43) aggregates (gain-of-function) are a specific marker of cognitive dysfunction^1^ and others have demonstrated gain-of-function roles in preclinical^13–15^ and clinical studies^16^.

Despite there being evidence for TDP-43 contributing to the pathogenesis of ALSFTSD through both gain- and loss-of-function mechanisms, there is little evidence examining the relative contribution, and temporal nature of these mechanisms in human tissue. Here, we have developed tools to examine both gain- and loss-of-function in a deeply clinically phenotyped cohort of *post-mortem* ALSFTSD cases. Our previously published^17–19^ cohort is composed of individuals stratified by clinical and pathological features across a range of disease subtypes (sporadic ALS, ALS-*SOD1* and ALS-*C9orf72*), in brain regions associated with cognitive function (executive, language and fluency), and cognitive symptom presentations (with or without cognitive deficits measured during life using the Edinburgh Cognitive ALS Screening tool). Using extra-motor brain regions has the advantage of profiling patients at different stages of the disease spectrum allowing us to probe the temporal nature of these mechanistic disease drivers (Figure 1). Using this temporally stratified cohort, we have developed tools to detect cryptic splicing events (single molecule *in situ* hybridization probes) and a broader range of TDP-43 aggregation events (a TDP-43 specific RNA aptamer^20^) to understand the relative contributions of gain and loss of TDP-43 function mechanisms in the pathogenesis of ALSFTSD.

**Figure 1.**
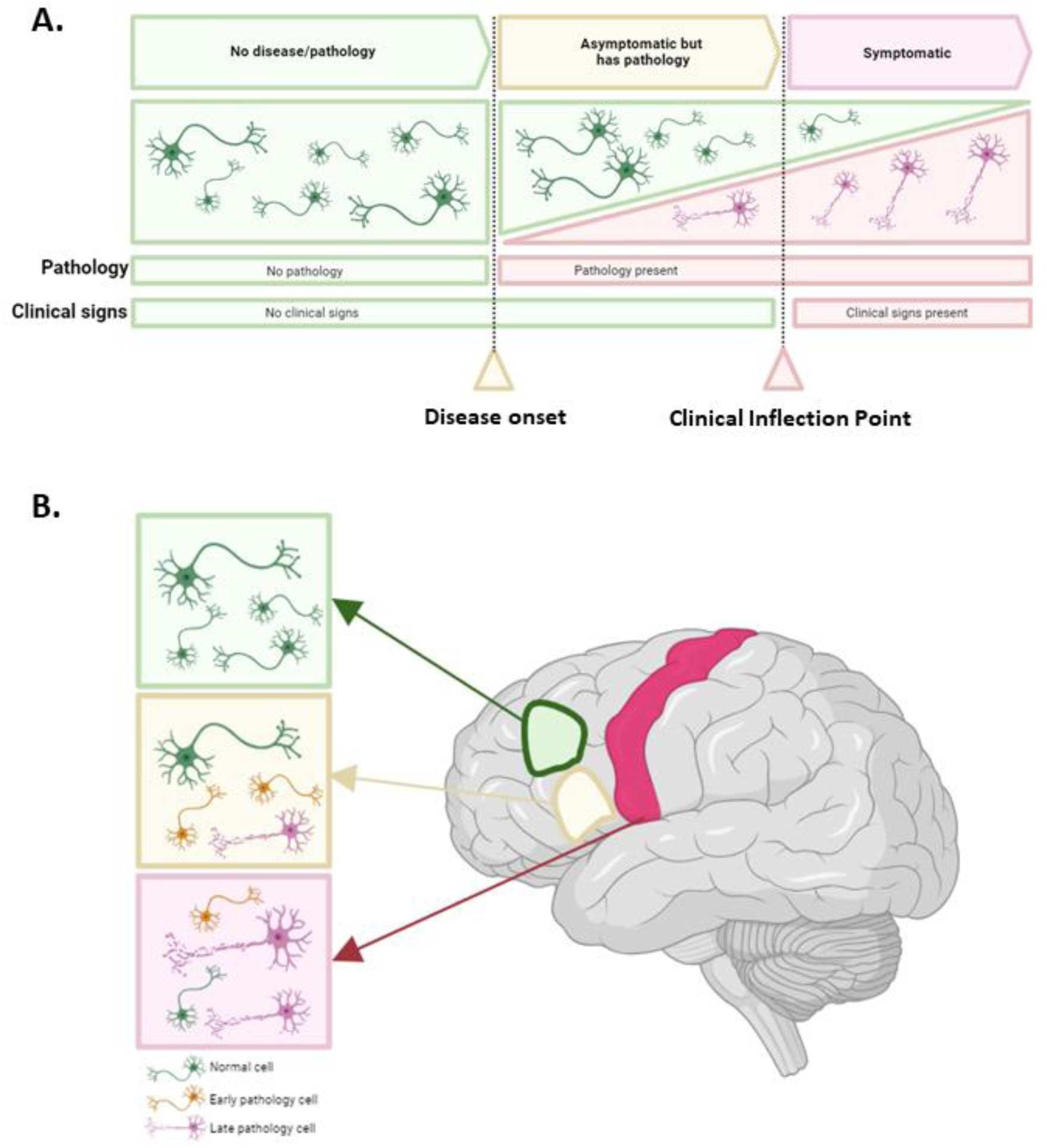
Schematic describing the temporal pathological changes seen in clinical progression. **A.** Diagram detailing the progression from disease and symptom-free (green), through asymptomatic pathology (yellow), to symptomatic pathology (red). **B**. Diagram illustrating that by examining distinct brain regions (BA46 in green; BA44 in yellow; and BA4 in red) with differential clinical involment (measured during life), it is possible to visualize temporal pathological stages ranging from no evidence of pathology or symptoms (green), through presence of pathology but no symptoms (yellow), to presence of pathology and symptoms (red). Noting that at any disease stage stage there may be a combination of affected and unaffected cells at different stages of the pathological process, even at *post-mortem*.

## Methods

### Case identification and cognitive profiling

Tissue was obtained from the Medical Research Council (MRC) Edinburgh Brain Bank (Table 1). All *post-mortem* tissue was collected with ethics approval from East of Scotland Research Ethics Service (16/ES/0084) in line with the Human Tissue (Scotland) Act (2006). Use of *post-mortem* tissue for studies was reviewed and approved by the Edinburgh Brain Bank ethics committee and the Academic and Clinical Central Office for Research and Development (ACCORD) medical research ethics committee (AMREC). Clinical data were collected as part of the Scottish Motor Neurone Disease Register (SMNDR) and Care Audit Research and Evaluation for Motor Neurone Disease (CARE-MND) platform, with ethics approval from Scotland A Research Ethics Committee (10/MRE00/78 and 15/SS/0216) and have been published previously^17–19^. Donors underwent neuropsychological testing during life with the Edinburgh Cognitive and Behavioural ALS Screen (ECAS) and all patients consented to the use of their data.

**Table 1.** Clinically stratified cohort of ALS cases and controls. Table showing the breakdown of cases and controls selected to represent each of the clinically stratified groups. Groups include cases with two brain regions (BA44 – language and fluency and BA46 – executive function) for which we have clinical data taken during life. Groups are: controls with no evidence of a proteinopathy (non-ALS disease controls; i.e. no TDP-43 pathology and no dementia), disease controls (no TDP-43 pathology but have ALS, e.g. SOD1), ALS concordant (TDP-43 pathology and cognitive dysfunction in that brain region) and ALS discordant cases (TDP-43 pathology with no evidence of cognitive dysfunction in that brain region).

| BA44 (Language & Fluency) |  |  |  | BA46 (Executive) |  |  |  |
| --- | --- | --- | --- | --- | --- | --- | --- |
| Case ID | Cognitive deficit | TDP – Pathology | Case type | Case ID | Cognitive deficit | TDP – Pathology | Case type |
| SD026/12 | Yes | Yes | Concordant | SD028/12 | Yes | Yes | Concordant |
| SD049/15 | Yes | Yes |  | SD047/13 | Yes | Yes |  |
| SD032/12 | Yes | Yes |  | SD032/12 | Yes | Yes |  |
| SD008/13 | No | Yes | Discordant | SD006/17 | No | Yes | Discordant |
| SD033/15 | No | Yes |  | SD008/13 | No | Yes |  |
| SD004/16 | No | Yes |  | SD033/15 | No | Yes |  |
| SD012/13 | No | No | Disease control | SD012/13 | No | No | Disease control |
| SD014/16 | No | No |  | SD014/16 | No | No |  |
| SD015/16 | No | No |  | SD015/16 | No | No |  |
| SD032/16 | Control case | No | Non-disease control | SD032/16 | Control case | No | Non-disease control |
| SD030/18 | Control case | No |  | SD030/18 | Control case | No |  |
| SD011/18 | Control case | No |  | SD011/18 | Control case | No |  |

### The clinically diverse patient cohort

To build a suitable patient cohort for our study, we set out to establish numerically balanced groups of four “case types” for comparison: These were 1) “Concordant” cases, where cognitive deficits had been identified along with pTDP-43 brain pathology, 2) “Discordant” cases, where pTDP-43 brain pathology had been identified but not cognitive deficits, 3) “Disease control” cases, constituting ALS patients with identified ALS-associated mutations (*SOD1*) but where no brain pTDP-43 pathology or cognitive deficit was identified, and 4) “Non-disease control” cases, representing controls with no diagnosis of ALS, no proteinopathy at *post-mortem* and no cognitive deficits in life (Table 1). For each of two cognitive brain regions (BA44 associated with language and fluency, and BA46 associated with executive function) for which TDP-43 brain pathology had been investigated, we identified three individuals for each of the four case types above. For both cognitive brain regions, cognition measured by the Edinburgh Cognitive ALS Screening tool and brain pTDP-43 pathology information was available and published previously^17^.

### Immunohistochemistry and BaseScope^TM^ in situ hybridisation

Formalin-fixed, paraffin-embedded (FFPE) tissue was cut on a Leica microtome into 4 μm thick serial sections that were collected on Superfrost (ThermoFisher Scientific) microscope slides. Sections were baked overnight at 40°C before staining. Sections were dewaxed using successive xylene washes, followed by alcohol hydration and treatment with picric acid to minimise formalin pigment. For pTDP-43 protein staining, antigen retrieval was carried out in citric acid buffer (pH 6) in a pressure cooker for 30 min, after which immunostaining was performed using the Novolink Polymer detection system (Leica Biosystems, Newcastle, UK) with a 2B Scientific (Oxfordshire, UK) anti-phospho(409–410)-TDP-43 antibody at a 1 in 4000 dilution and a Novus stathmin-2 antibody (St. Louis, USA) at a 1:500 dilution with no antigen retrieval step.

For validation of *STMN-2* expression, two BaseScope^TM^ probes were designed, one that targets *STMN-2* downstream of exon 2 to detect normal *STMN-2* (*STMN-2(N);* Catalogue number 1048241-C1), and one that targets the *STMN-2* cryptic exon (*STMN-2(CE);* Catalogue number 1048231-C1). The BaseScope^TM^ protocol was performed as we have published previously with no additional modifications^18,19^. Slides were counterstained using haematoxylin and blued with lithium carbonate. *STMN-2* expression was quantified manually by a pathologist blinded to clinical and phenotypic data. Manual grading was performed by counting the number of transcripts per cell in 20 cells in each of three 40x fields of view per section. Whole tissue sections were scanned with brightfield illumination at 40x magnification using a Hamamatsu NanoZoomer XR. Using NDP.view2 viewing software (Hamamatsu), regions of interest (ROIs) were taken from key regions for quantification as described below.

### Immunohistochemistry modifications for RNA aptamer staining

Tissue was prepared in the same way for IHC as listed above. Following deparaffinisation, rehydration, and antigen retrieval, slides were incubated with peroxidase block for 30 minutes followed by 5-minute wash step with TBS. Avidin and biotin blocking steps were then carried out using a biotin blocking kit (ab64212) as per the manufacturer’s guidelines followed by a 5-minute TBS wash step and a 5-minute wash step with milli-Q water. 156nM of aptamer (TDP-43^Apt^ CGGUGUUGCU with a 3’ Biotin-TEG modification, ATDBio, Southampton, UK) prepared in Milli-Q water was applied to the tissue and incubated for 3 hours at 4°C followed by incubation with 4% PFA overnight at 4°C. A 5-minute wash with Milli-Q water then preceded incubation with anti-biotin HRP antibody (ab6651) diluted 1 in 200 in milli-Q water for 30 minutes followed by a 5-minute wash step with Milli-Q water and incubation with DAB for 5 minutes. Slides were then counterstained, dehydrated and cleared as detailed above. For double staining, the entire BaseScope^TM^ staining protocol was implemented and following the application of red chromogen, slides were washed with TBS for 5 mins and then taken straight to the avidin/biotin blocking steps of the aptamer staining protocol.

### Immunofluorescence staining

Tissue was prepared in the same way for IHC as listed above. Following deparaffinisation, rehydration, and antigen retrieval, slides were permeabilised in PBS + 0.3% Triton-X for 15mins and then rinsed with PBS. Autofluorescence quenching was then performed by incubating slides for 2min in 70% ethanol, followed by 0.1% Sudan Black in 70% ethanol (20mins) then a further 2 mins in 70% ethanol. Slides were then washed in PBS followed by a 5 min incubation with TrueView (prepared as 1:1:1 components A, B and C). Slides were then washed for 5 mins in PBS and blocked for 1hr in blocking buffer (PBS supplemented with 1% goat serum and 0.1mg/ml salmon sperm DNA). Slides were then incubated overnight at 4°C with 1.2 μg/mL pTDP-43 primary antibody (Proteintech, 22309-1-AP) prepared in blocking buffer. Three 5 min washes with PBS preceded a 2-hour incubation at room temperature with 4 μg/mL Alexa Fluor 647-conjugated secondary antibody (ThermoFisher A21244) prepared in blocking buffer. Three 5 min wash steps with PBS were then preformed followed by incubation with 1.2 μg/mL C-terminal TDP-43 Antibody with TDP-43^Apt^ (CGGUGUUGCU with a 3’ Biotin-TEG modification, ATDBio, Southampton, UK) [0.67 μM] prepared in blocking buffer for 4hr at RT in the dark. A wash step was performed with blocking buffer with aptamer E2108 [0.67 μM i.e. 1:50]. Slides were then incubated with 4% PFA in PBS (30min), washed with PBS and then incubated with DAPI 1:10,000 (10min), washed with PBS and mounted with Vectashield anti-fade. Three 5 min wash steps with PBS were then preformed followed by 4hr covered incubation at room temperature with 1.2 μg/mL CoraLite Plus 488-conjugated C-terminal TDP-43 antibody (Proteintech, CL488-12892) and 0.67 μM TDP-43^Apt^ (CGGUGUUGCU with a 3’ Atto 590 modification, ATDBio, Southampton, UK) prepared in blocking buffer. A wash step was performed with blocking buffer supplemented with 0.67 μM Atto 590-conjugated TDP^APT^. Slides were then incubated with 4% PFA in PBS (30min), washed with PBS and then incubated with DAPI 1:10,000 (10min), washed with PBS and mounted with Vectashield anti-fade. Sections were imaged using a Zeiss AxioScan Z1 slide scanner with identical acquisition settings for all images.

### Quantitative digital and manual pathology analysis

Ten (400 x 400 pixel) regions of interest (ROIs) were taken from each whole slide scanned image. Neuropil, neuronal nuclei, and neuronal cytoplasm were manually segmented using QuPath software^21^. Mean DAB intensity for neuropil, neuronal nuclei and neuronal cytoplasm were calculated and exported for subsequent analysis. DoG superpixel segmentation analysis was then carried out on the neuropil, neuronal nuclei, and neuronal cytoplasm separately to quantify aptamer-positive foci. Here, compartments were split into superpixels generated from pixels with similar intensities and textures for further classification, and each superpixel was classified as positive or negative for aptamer dependent on pre-set DAB intensity thresholds. The area of positive superpixels and total area were then exported. Weighted mean DAB intensity and weighted mean area of positive superpixels (where total area was used to weight) were then calculated per case (10 ROIs) to prevent pseudo replication. ROIs were then blinded and manually rated for (i) number of neurons with rod features (Nrods), (ii) more than one rod feature (N>1rod+1), (iii) nuclear aggregation (Nnuclearaggregation+1), (iv) membrane pathology (Nmembranepath+1), and (v) punctate cytoplasmic stain (Ncytoplasmicpuncta+1). Blinded ROIs were also manually rated for the numbers of glia with (i) nuclear pathology and (ii) with cytoplasmic pathology. The product score for all neuronal features was calculated as

((Nrods+1)*(N>1rod+1)*(Nnuclearaggregation+1)*(Nmembranepath+1)*(Ncytoplasmicpuncta+1))

Weighted mean neuronal features (where total number of neurons was used to weight) and weighted mean glial features (where total number of glia were used to weight) were calculated per case (10 ROIs) to prevent pseudoreplication. Data were visualized using RStudio with the “ggplot2” package^22^. ANOVA was used to determine differences between controls, discordant and concordant groups. Pearsons or Spearman’s correlation tests were used to determine correlations between *STMN-2* counts and aptamer scores.

## Results

### STMN-2 cryptic splicing events, but not pTDP-43 pathology, distinguish between distinct clinical phenotypes

*STMN-2* cryptic splicing pathology has been shown to be an accurate molecular marker of TDP-43 loss-of-function in ALS-TDP^8^. Here, we set out to develop tissue-based detection probes to identify, using *in situ* hybridization, *STMN-2* cryptic splicing events and to understand their temporal relationship with TDP-43 pathology (detected by the pTDP-43 antibody) and clinical phenotype (by looking at cases covering a spectrum of clinical presentations). Clinical presentations included non-neurological controls and disease controls that had no evidence of pTDP-43 pathology or cognitive dysfunction, as well as cases that did have pTDP-43 pathology, segregated into i) concordant cases, which had cognitive dysfunction, and ii) discordant cases, which did not have cognitive dysfunction. Using this clinically and molecularly defined cohort, we first demonstrate normal STMN-2 protein expression using immunohistochemistry (Figure 2A) in non-neurological and disease controls. The characteristic immunophenotype of STMN-2 protein distribution is that of a marginated crisp cytoplasmic stain forming a complete ring around the neuronal cell. This immunostaining corresponds to the presence of a BaseScope^TM^ *in situ* hybridisation signal of abundant normal *STMN-2 (STMN-2(N))* mRNA transcripts and the absence of *STMN-2* cryptic exon transcripts (*STMN-2(CE)*) (Figure 2A). In concordant cases, we observed abundant *STMN-2(CE)* cryptic exon transcripts, and an absence of full length *STMN-2(N)* mRNA, a finding that corresponded to the absence of STMN-2 protein immunoreactivity in neurons (Figure 2A), illustrating that only the expression of the *STMN-2(N)* mRNA transcript results in normal STMN-2 protein expression. Interestingly, in discordant cases, both *STMN-2(CE)* and *STMN-2(N)* could be visualized within spatially distinct regions on serial sections. Quantification of the expression of each of these transcripts in brain regions BA44 and BA46, performed blinded to clinical and demographic information, demonstrates that their abundance remains consistent between phenotypically similar cases and that *STMN-2(CE)* expression is consistently present in cases with pTDP-43 pathology, but that *STMN-2(N)* is substantially reduced or completely absent only when cases also show clinical manifestation of that pathology (i.e., concordant cases; Figure 2B).

**Figure 2.**
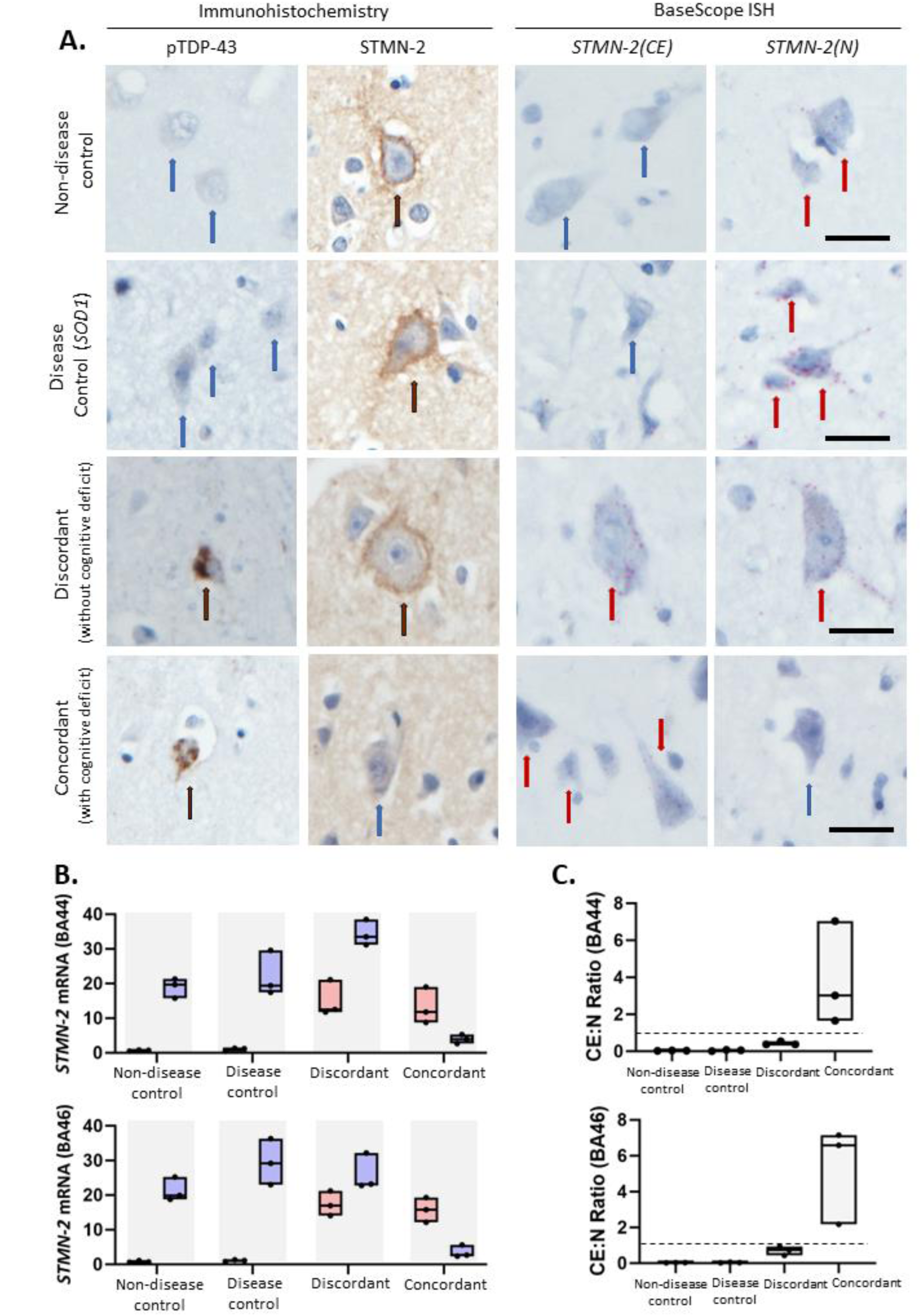
STMN-2 cryptic splicing events, but not pTDP-43 pathology, distinguish between distinct clinical phenotypes. **A.** Representative photomicrographs taken at 40x magnification demonstrating neuronal phospho-TDP-43 pathology and Stathmin-2 (protein) staining using immunohistochemistry (left two panels), as well as *Stathmin-2* cryptic exon (CE) and *Stathmin-2* normal mRNA transcripts using BaseScope^TM^ *in situ* hybridization (right two panels). Blue arrows indicate absence of immunoreactivity, black arrows indicate positive staining on immunohistochemistry (protein) and red arrow indicate postitive staining on *in situ* hybridisation (mRNA). Each red dot represents a single mRNA transcript. Scale bar = 50 µm. **B.** Box plots demonstrating phenotypically conserved variation in *Stathmin-2* cryptic exon (CE) and *Stathmin-2* normal transcripts per cell in the two brain regions examined (BA44 and BA46). Each bar of the box plot represents data from three cases, with medians (horizontal line on box), spread and skewness derived from counts of transcripts per cell from 20 cells in each of three randomly allocated regions of interest, boxes represent min and max values. Graphs demonstrate counts to be consistent between clinically segregated groups with cases expressing more cryptic exon than controls and concordant cases demonstrating the most striking loss of normal *Stathmin-2* expression. **C**. Box plots showing the ratio of cryptic exon to normal counts. Each bar of the box plot represents data from three cases, with medians (horizontal line on box), spread and skewness derived from ratio of counts shown in B, boxes represent min and max values. Horizontal dotted line at *y* = 1, shows that only concordant (clinically manifesting) cases have a ratio of >1.

Notably, general patterns of *STMN-2(CE)* and *STMN-2(N)* expression were similar for both language- and fluency-associated region BA44, as well as executive function-associated BA46 region, across all four case types (Figure 2B). Specifically, for both brain regions, *STMN-2(CE)* expression was very low, but not completely absent, in *SOD1* (disease controls) and absent in non-disease controls, but high in concordant and discordant cases (both with pTDP-43 pathology). Normal *STMN-2(N)* expression in both regions was high in disease controls and non-disease controls, low for concordant cases, but moderate to high in discordant cases (Figure 2B). Therefore, whilst the presence of *STMN-2(CE)* does appear to be an early pathological feature, preceding clinical symptoms, a combination of the presence of *STMN-2(CE)* and the loss of *STMN-2(N)* expression, calculated as a ratio (Ratio of >1 is only seen in concordant cases; Figure 2B), is better predictor of clinical phenotype than cryptic exon presence alone.

### TDP-43^APT^ identifies broader range of aggregation events compared to classical antibody approaches

Using this molecularly and clinically phenotyped tissue cohort, we next explored the role of toxic gain-of-function. As pTDP-43 staining was not able to distinguish between differentially clinically stratified cases, we developed a modified staining technique to utilize our recently developed TDP-43 specific RNA aptamer^20^. Taking advantage of the ability to easily modify RNA aptamers, we first biotinylated TDP-43^APT^ for immunohistochemical staining. We subsequently developed a staining technique, requiring a post-staining fixation step (Supplementary Figure 2), that achieves specific staining of TDP-43 pathology (Figure 3A; left panel). Using this approach, we identified both neuronal and glial pathology in both the cytoplasm and nucleus of affected cells in ALS cases, but not controls (Figure 3B). Immunofluorescent staining was performed using three markers of TDP-43 pathology: (i) the pTDP-43 antibody (purple), (ii) the C-terminal TDP-43 antibody (green), and (iii) TDP-43^APT^ aptamer (red). TDP-43^APT^ was able to detect aggregated TDP-43 in the cytoplasm of affected cells (cell labelled 1) as well as nuclear accumulation (cell labelled 2), that would ordinarily have been obscured by “normal” (i.e., functional, non-phosphorylated) C-terminal TDP-43 antibody staining (Figure 3B). Additional examples of TDP-43^APT^ binding to diverse aggregate morphologies (including cytoplasmic and nuclear features) are detailed in Supplementary Figure 3. The pathological features that the RNA aptamer can detect include nuclear membrane staining as well as nucleolar decoration, nuclear puncta, and nuclear rods (occasionally multiple nuclear rods) (Figure 3C). Cytoplasmic neuronal aggregates can also be identified (Figure 3D) as well as glial pathology (both nuclear and cytoplasmic (Figure 3E)).

**Figure 3.**
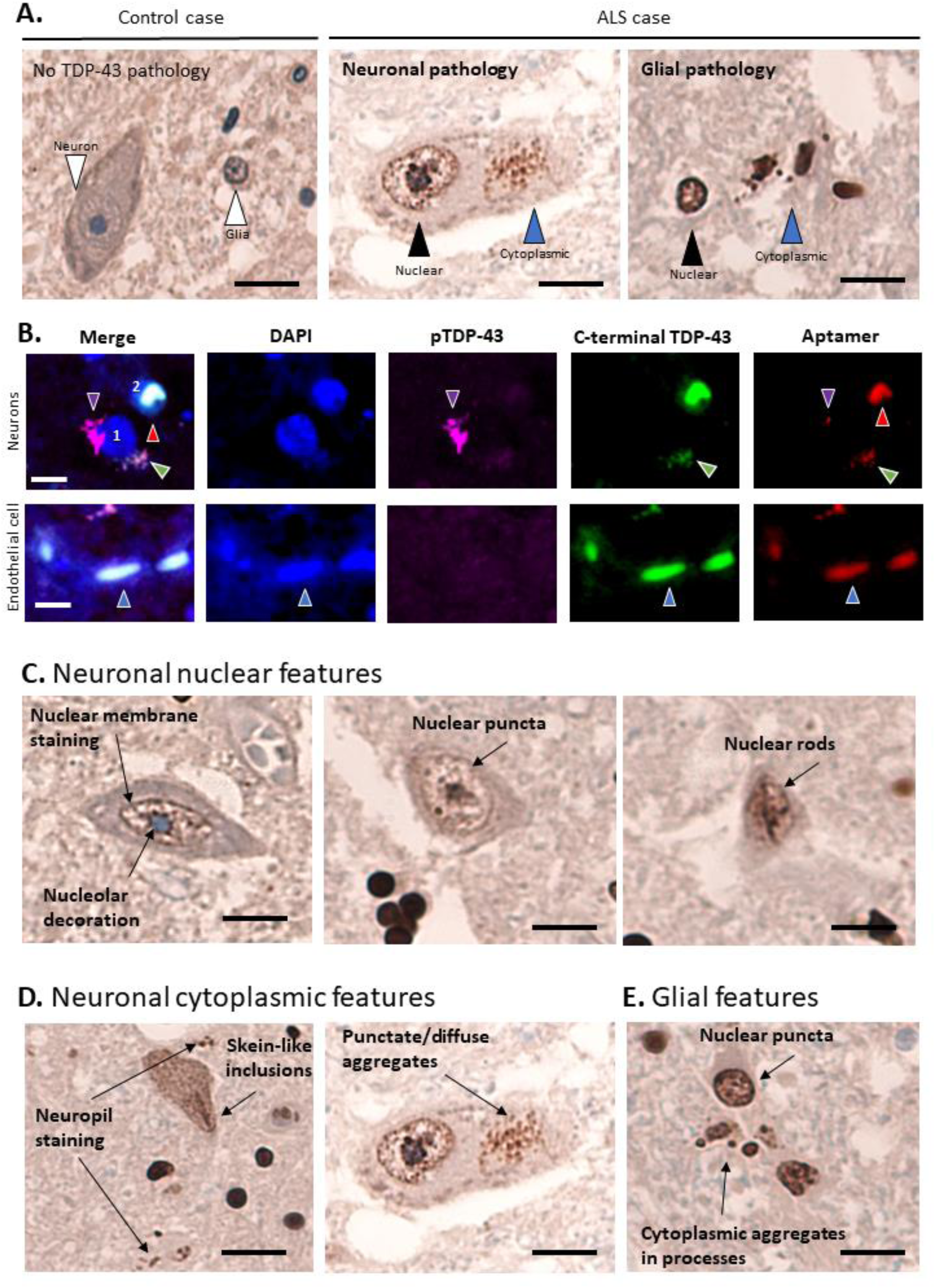
TDP-43^APT^ identifies broader range of aggregation events compared to classical antibody approaches. **A.** Representative photomicrographs taken at 40x magnification demonstrating the staining pattern of neuronal and glial cells with a biotiylated RNA aptamer targeting TDP-43 (TDP43^APT^). Controls (left image) show no evidence of immunoreactivity, and sALS cases (centre and right images) show neuronal nuclear (black arrowhead) and cytoplasmic pathology (blue arrowhead; centre image) and glial nuclear (black arrowhead) and cytoplasmic pathology (blue arrow head; right image). Scale bar is 20 µm **B.** Representative photomicrographs taken at 20x magnification demonstrating immunofluorescent staining with a pTDP-43 antibody (purple), a c-terminal TDP-43 antibody (green) and TDP-43^APT^ (red). Scale bar is 5 µm and nuclei are stained with DAPI (blue). Images show two neurons (nuclei labeled 1 and 2) affected by TDP-43 pathology. Cell 1 shows two cytoplasmic aggregates, one aggregate co-stains with the pTDP-43 antibody and TDP-43^APT^ (purple arrowhead) and the other aggregate co-stains with the c-terminal antibody and TDP-43^APT^ (green arrowhead). The cell labelled 2 shows a nuclear aggregate (red arrowhead), identified by the TDP-43^APT^, but which is obscured by the “normal” (i.e., functional, non-phosphorylated) C-terminal antibody staining. Endothelial cells (lower panel) that are not involved by TDP-43 pathology show only diffuse, non-aggregated C-terminal antibody and TDP-43^APT^ staining (blue arrowheads) and show no immunoreactivity for pTDP-43. **C-E.** Representative photomicrographs taken at 40x magnification demonstrating DAB immunostaining for TDP-43^APT^ highlighting neuronal nuclear features (**C**), neuronal cytoplasmic features (**D**) and glial TDP-43 pathology (**E**). Scale bar = 20 µm.

### TDP-43^APT^ pathology correlates with cryptic exon splicing events

We next sought to understand the relationship between TDP-43 loss-of-function (using our *STMN-2* ISH probes) and TDP-43 gain-of-function, using our TDP-43^APT^ staining protocol. To do this, we used tissue from a discordant case where there were distinct regions that had both preservation of *STMN-2* normal expression, as well as loss within the same section. Using this approach, it was possible to visualize examples of cells where *STMN-2(N)* mRNA transcripts are detected and where there is no evidence of TDP-43^APT^ pathology. Correspondingly, where there are cells with abundant TDP-43^APT^ pathology, we see a reciprocal complete loss of *STMN-2(N)* mRNA transcripts (Figure 4A, upper panel). We also demonstrate the reciprocal findings for *STMN-2(CE)* mRNA transcripts and aptamer features (Figure 4A, lower panel). Indeed, cryptic exons appear to be present even with very mild (nuclear only) TDP-43^APT^ pathology in the absences of significant cytoplasmic pathology (Figure 4C). These features were quantified using a combination of pathological features derived from both digital and blinded manual grading of the cohort (Supplementary Figure 3), we observe a significant correlation between *STMN-2* splicing pathology and TDP-43 pathology in the form of nuclear (*R* = 0.82; *p* = 0.0068) and cytoplasmic puncta (*R* = 0.73; *p* = 0.021) (Figure 4C). A summary for all pathological features is included in Figure 4D. Importantly, this analysis was performed on individually stained slides to prevent confounding effects of steric hindrance within the co-stained slides.

**Figure 4.**
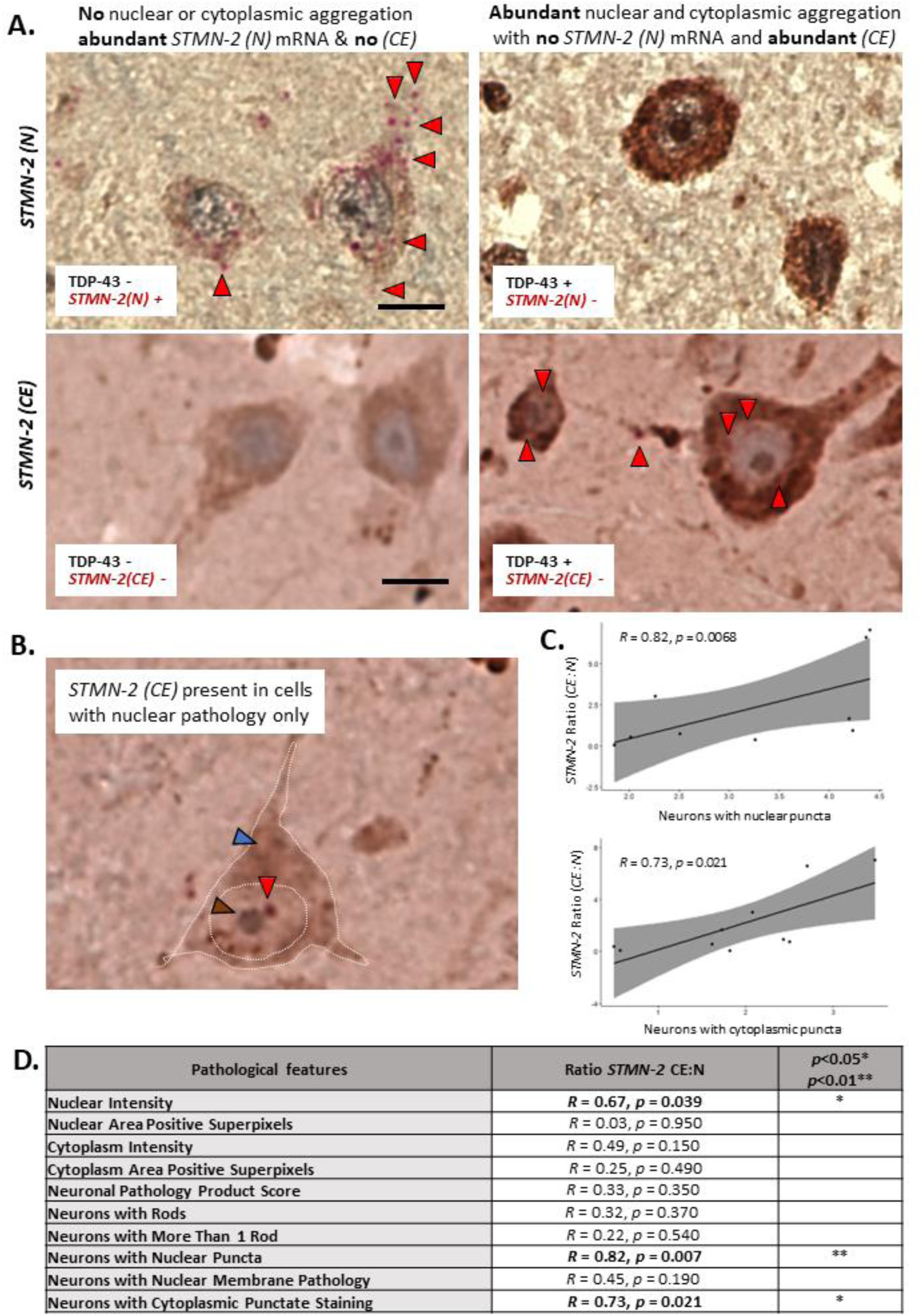
TDP-43 pathology detected by RNA aptamer correlates with molecular phenotype. **A.** Representative photomicrographs taken at 40x magnification demonstrating dual DAB immunohistochemical staining for TDP-43^APT^ and *in situ* hybridization with BaseScope^TM^ to detect *STMN-2* normal (N) and cryptic exon (CE) mRNA transcripts (individual red dots are single mRNA transcripts of *STMN-2*). Images are taken from distinct regions of a case with a discordant clinical phenotype (i.e. TDP-43 pathology present but no evidence of clinical manifestation) showing that in regions where there is no evidence of TDP-43^APT^ pathology there is ample normal *STMN-2* mRNA expression (red arrowheads; top left image) and no cryptic exons present (lower left image). However, in regions where there is abundant nuclear and cytoplamsic aggregation seen with TDP-43^APT^ staining (right images), there is a coincident loss of normal *STMN-2* expression (upper right image) and cryptic exons can be seen (red arrowheads; lower right image). Scale bar = 20 µm. **B.** Image demonstrating the presence of cryptic exons (red arrowheads) in a neuron with nuclear pathology (brown arrowhead) in the absence of substantial cytoplasmic pathology (blue arrowhead). **C.** Graph demonstrating the correlation between *STMN-2* splicing pathology and TDP-43^APT^ pathology with respect to nuclear (left graph) and cytoplasmic (right graph) TDP-43 pathology. **D.** Summary of all comparisons made between TDP-43^APT^ pathology (pathological features) and *STMN-2* cryptic splicing events.

### Nuclear TDP-43^APT^ pathology is an early event that coincides with TDP-43 loss-of-function and precedes clinical symptom onset

Having demonstrated that TDP-43 loss-of-function, measured by *STMN-2* cryptic splicing events, was associated with clinical phenotype and TDP-43^APT^ pathology, we next wanted to understand the distribution of TDP-43^APT^ features in our cohort and how distinct subcellular pathologies might relate to the temporal progression of clinical phenotype. We noted both cytoplasmic and nuclear pathological features in concordant, clinically manifesting regions, but more variable and sparser pathology, which was predominantly nuclear in discordant (pre-symptomatic) regions (Figure 5A). This was quantified using a combination of digital and blinded manual assessment of pathological features (Supplementary Figure 3), demonstrating that the extent of neuronal and glial pathology was associated with clinical phenotype, with statistically significant differences between cases and controls as well as between concordant and discordant cases (Figure 5B). These differences were largely driven by nuclear, but not cytoplasmic features, the extent of which appear to determine clinical phenotype (Figure 5C,D). All statistical associations between pathological features and clinical phenotype are summarised in Table 2. These data support the hypothesis that TDP-43^APT^ nuclear staining, is an early event that occurs, along with *STMN-2* cryptic exon emergence, prior to the clinical inflection point between pre-symptomatic and symptomatic states.

**Figure 5.**
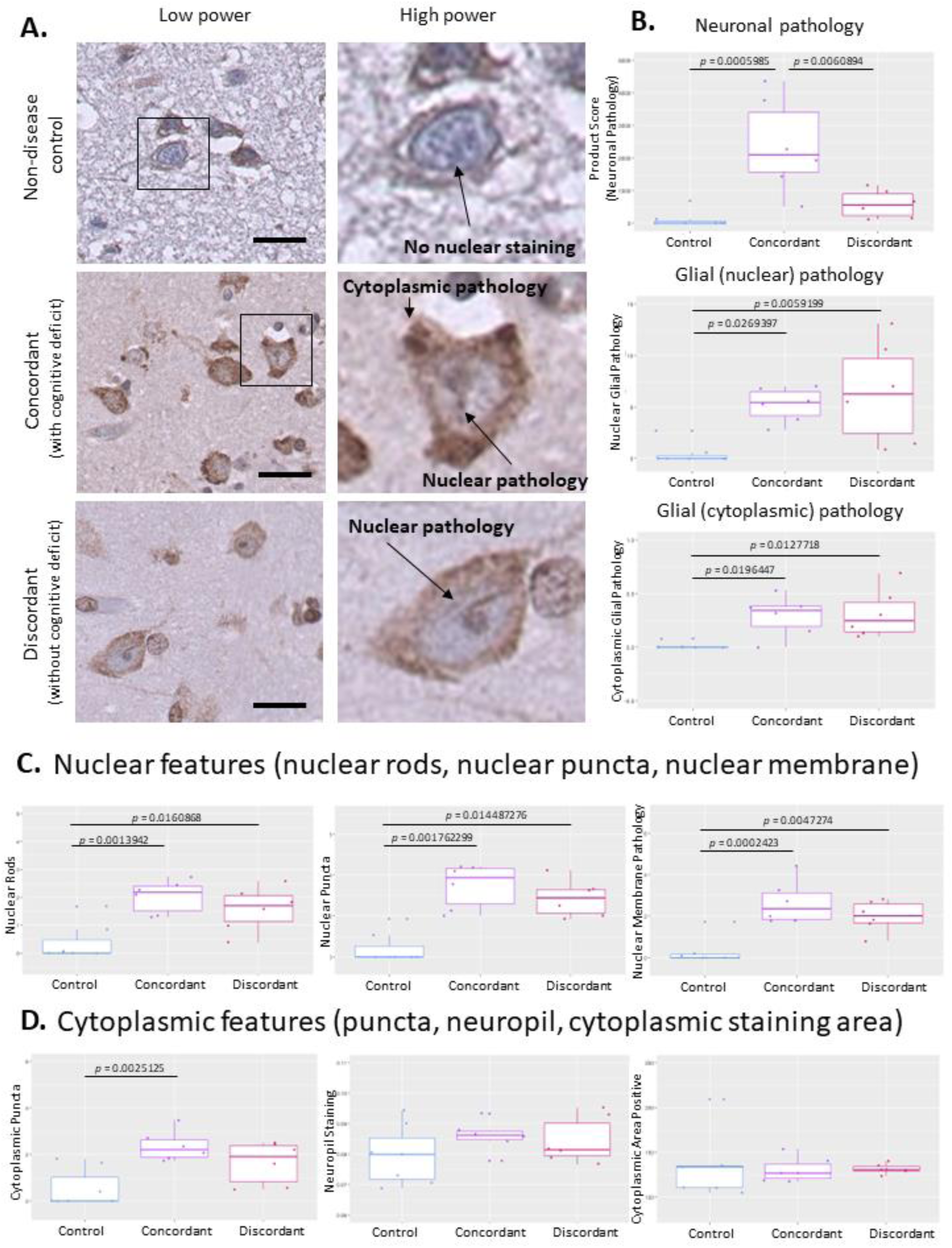
Nuclear TDP-43^APT^ pathology is an early event that coincides with TDP-43 loss-of-function and precedes clinical symptom onset. **A.** Representative photomicrographs taken at 40x magnification demonstrating the staining pattern of TDP-43^APT^ in cases that have been clinically stratified in to non-neurological, concordant and discordant cases. Images on the left show the low power region that the digital zoom has focused on in the right panel. Nuclear and cytoplasmic features have been annotated on the images. Scale bar = 50 µm. **B.** Graphs demonstrating a product score of all neuronal pathologies (upper graph; ANOVA corrected for multiple comparisons shows statistically significant difference between groups, *p* = 0.000552), glial nuclear pathology (centre graph; ANOVA corrected for multiple comparisons shows statistically significant difference between groups, *p* = 0.00489), and glial cytoplasmic pathology (lower graph; ANOVA corrected for multiple comparisons shows statistically significant difference between groups, *p* = 0.0071), examined using digital and blinded manual assessment of images represented in A. **C.** Graphs demonstrating neuronal nuclear pathologies including the number of cells with more than one visible nuclear rod (left graph; ANOVA corrected for multiple comparisons shows statistically significant difference between groups, *p* = 0.00136), nuclear puncta (centre graph; ANOVA corrected for multiple comparisons shows statistically significant difference between groups, *p* = 0.001262), and nuclear membrane pathology (right graph; ANOVA corrected for multiple comparisons shows statistically significant difference between groups, *p* = 0.000238), examined using digital and blinded manual assessment of images represented in A. **D.** Graphs demonstrating neuronal cytoplasmic pathologies including cytoplasmic puncta (left graph; ANOVA corrected for multiple comparisons shows statistically significant difference between groups, *p* = 0.0025), neuropil staining (centre graph; ANOVA corrected for multiple comparisons shows statistically significant difference between groups, *p* = 0.964), and cytoplamsic staining area (right graph; ANOVA corrected for multiple comparisons shows statistically significant difference between groups, *p* = 0.4019), examined using digital and blinded manual assessment of images represented in A.

**Table 2.** Statistical associations between pathological features and clinical phenotype.

| Pathology | Case v Controls | Concordant v Controls | Discordant v Controls | Concordant v Discordant |
| --- | --- | --- | --- | --- |
| Neuropil Intensity |  |  |  |  |
| Neuropil Area Positive Superpixels |  |  |  |  |
| Nuclear Intensity |  |  |  |  |
| Nuclear Area Positive Superpixels | * | * |  |  |
| Cytoplasm Intensity |  |  |  |  |
| Cytoplasm Area Positive Superpixels |  |  |  |  |
| Neuronal Pathology Product Score | ** | *** |  | ** |
| Neurons with Rods | *** | ** | ** |  |
| Neurons with More Than 1 Rod | *** | ** | * |  |
| Neurons with Nuclear Puncta | *** | ** | * |  |
| Neurons with Nuclear Membrane Pathology | *** | *** | ** |  |
| Neurons with Cytoplasmic Punctate Staining | ** | ** |  |  |
| Glia with Nuclear Pathology | * | * | ** |  |
| Glia with Cytoplasmic Pathology | ** | * | * |  |

Table summarising all statistical comparisons made between clinically stratified cases for all pathological features examined. Differences between cases and controls and controls and concordant cases were driven primarily by the combination of nuclear and cytoplasmic pathology. However differences between discordant and control cases were primarily driven by neuronal nuclear (but not cytoplamsic) features and glial pathology. These associations imply that nuclear features are an early sign of disease, but that widespread cytoplasmic pathology is required to show clinical manifestations.

## Discussion

In this study, we aimed to understand the relative contributions of gain- and loss-of-function mechanisms underpinning TDP-43 pathology in ALSFTSD. To do this, we developed tools to probe gain-of-function (using TDP-43^APT^ to identify aggregation events not readily visualised using classical antibody staining) and loss-of-function (*STMN-2* cryptic exon signatures) in a temporally, clinically stratified *post-mortem* tissue cohort. We identify a temporal loss-of-function of TDP-43 starting with individuals characterized by differentially affected pools of neurons, some with evidence of cryptic splicing events and some with normal *STMN-2* splicing, who had not yet begun to develop symptoms of cognitive decline and ending with clinically manifesting individuals who had a complete loss of *STMN-2* expression. We have shown previously^17^ that in non-demented patients, TDP-43 pathology across a wide range of brain regions is a robust, specific marker of ALS across a range of cognitive dysfunction. Importantly, however, the sensitivity of TDP-43 pathology as a marker of cognitive dysfunction is poor - 17.7 % for executive function (3.8-43.4, 95% CI), 47.1% for language function (23.0-72.2, 95% CI), and for 26.7% letter fluency (6.80-55.1, 95% CI)^17^. Indeed, Prudencio *et al.* ^8^ found truncated *STMN-2* to be a highly specific biomarker for FTLD-TDP, as *STMN-2* was absent in brain tissues of patients with FTLD-FUS, FTLD-tau and in controls, and was absent in spinal cords of patients with ALS-SOD1 and in controls. However, *STMN-2* seems a less sensitive marker in the frontal cortex (82.4%) and in the spinal cord (62.2%) of patients with FTLD-TDP. Although the *STMN-2* cryptic exon was less sensitive, it was still appreciably more sensitive than pTDP-43 at detecting clinical phenotype in these previous studies. In our cohort of cases, we consistently observed either loss of normal *STMN-2* transcripts or gain of *STMN-2* cryptic exons (or a combination of both), making the ratio of *STMN-2* cryptic to normal *STMN-2* the most robust measure of pathology in these cases (Figure 2C). We also observed that the extent to which these features were evident within cases, using this ratio advantage, was highly correlated with clinical phenotype, a finding that has also been seen in other cohorts where truncated *STMN-2* has been shown to be associated with earlier age of onset of ALS-FTD^8^. Therefore, combined with our data demonstrating that improved diagnostic accuracy can be achieved using the ratio of truncated to full-length *STMN-2*, this could provide additional insights into the likely development of cognitive symptoms in this population.

This temporal spectrum of molecular features of TDP-43 loss-of-function preceding the clinical inflection point (i.e., present prior to clinical manifestation in individuals/brain regions) also mapped on to TDP-43 pathology in distinct cell compartments. Using TDP-43^APT^, we identified early nuclear events which appeared to precede more extensive pathology involving both the nucleus and cytoplasm when individuals showed clinical signs in these brain regions. Indeed, nuclear TDP-43 pathology has been reported previously, with cells expressing GFP-tagged C-terminal fragments of TDP-43^23^ exhibiting irregularly shaped nuclei with deep invaginations of the nuclear membrane (imaged using electron microscopy), and with morphological features resembling the nuclear rods described here (see Figures 3 and 5). Furthermore, a recent study demonstrated differences between the morphology and ubiquitylation of cytoplasmic TDP-43 aggregates, where they observed premature and poorly ubiquitylated TDP-43 inclusions to be associated with high levels of nuclear TDP-43, whereas mature and well-ubiquitylated inclusions were associated with nuclear clearance of TDP-43^24^. Additionally, recent work characterising CHMP7 accumulation in sporadic and familial ALS which has been shown to lead to dysfunction of the nuclear pore, leading to nuclear TDP-43 dysfunction, is consistent with our findings of temporal progression from early nuclear to later cytoplasmic pathology^25^. Crucially, TDP-43^APT^ now provides the possibility to examine these pathological processes as it can detect a wider range of aggregation events, including nuclear pathology, compared to standard antibody approaches alone. This also raises the possibility of early intervention as these early nuclear events, in our cohort, are in the pre-symptomatic disease phase raising the possibility of early, pre-symptomatic targeting of nuclear pore pathology. The improved sensitivity and specificity of TDP-43^APT^ in detecting TDP-43 pathology, if adapted as a PET tracer or liquid biomarker, could aid in the identification of individuals for targeted early interventions.

It should also be noted that, as we do not yet know of any cryptic splicing targets for non-neuronal cells, we did not explore the role of glial pathology beyond profiling TDP-43 pathology with TDP-43^APT^. However, we do demonstrate intriguing glial TDP-43^APT^ pathology (Figure 5B). Notably, nuclear pathology occurs early and is more variable and not linked to clinical phenotype beyond being a sensitive measure of case vs. control. Further interrogation of these findings, including interpretation of the clinical and molecular context, is clearly warranted once specific cryptic splicing targets are discovered. Our own previous studies examining this stratified cohort have demonstrated cell-type specific differences for both (i) protein folding capacity including chaperone proteins (clusterin)^19^, and (ii) inflammatory markers like the NLRP3 inflammasome^18^, indicating that cell-type specific changes may well underpin differential susceptibilities to signs of clinical deterioration in these brain regions.

TDP-43^APT^, is designed to have optimal affinity to bind to the RNA recognition motifs of TDP-43. Crucially, clinical implementation of RNA aptamers has been limited due to the requirement for these molecules to fold into appropriate secondary structures^26^. However, as this TDP-43 aptamer is designed to maintain a single-stranded structure, it is not subject to these limitations. Here we have adapted this aptamer for use with immunohistochemical staining using a biotin tag, demonstrating high specificity for pathological TDP-43, with cells unaffected by TDP-43 pathology (e.g., endothelial cells in Figure 3B) showing no immunoreactivity. Indeed, a range of other molecular tags are also possible, including fluorophores as we have published previously^20^. Given the unprecedented sensitivity and specificity of TDP-43^APT^ for disease-related pathology that we have demonstrated in this study, there is also the possibility that this aptamer could be tagged and used as a radiolabel for PET imaging or as a contrast agent for MRI^27,28^, as other aptamers have been used to detect amyloid plaques in Alzheimer’s disease. Other applications include using TDP-43^APT^ to detect pathology in peripheral samples such as CSF and serum. The ability to differentiate early pre-symptomatic disease states and the relative ease and cost-effectiveness of developing and synthesising RNA probes compared to antibody generation, combined with the flexibility to incorporate a diverse array of functional tags are clearly advantageous properties compared to conventional antibody approaches. Further studies are clearly warranted to understand how the improved detection accuracy afforded by TDP-43^APT^ could improve how pathology could be detected in a clinical setting.

Taken together, our findings demonstrate the temporal progression of TDP-43 pathology in non-motor brain regions in ALS and support the use of TDP-43^APT^ and *STMN-2* cryptic splicing events as informative readouts for both motor and non-motor clinical manifestations. Through utilisation of a TDP-43 aptamer, we uncover and describe a wide-range of TDP-43 aggregation events encompassing early nuclear-only pathology through to later stage extensive cytoplasmic and nuclear pathology. The pathological markers of loss-of-function, notably early nuclear pathology, as well as the *STMN-2* cryptic splicing pathology appear to manifest prior to the onset of clinical symptoms, indicating that these markers could provide targets for accurate early diagnosis but are not required for clinical manifestation. Our findings raise the possibility that the presence of *STMN-2* cryptic exons in biofluids could be used as a marker of early disease prior to symptom onset. Furthermore, the ratio of CE:N *STMN-2* could provide a reliable method to predict phenoconversion as ultimately the combination of cryptic exon presence as well as loss of normal *STMN-2* is the best predictor, in our data, of clinical phenotype and that extensive TDP-43 cytoplasmic aggregation (gain-of-function) is required for clinical manifestation.

## Acknowledgements

The research leading to this manuscript has been supported by (i) a Target ALS foundation grant to JMG, MHH, GGT, EZ and NS and employing MG and FMW BB-2022-C4-L2; (ii) an NIH grant to JG and MHH, employing HS and FR R01NS127186; (iii) the European Research Council (RIBOMYLOME_309545 and ASTRA_855923) to GGT; and (iv) an MND Association Lady Edith Wolfson Junior Non-Clinical Fellowship to RS Saleeb/Oct22/980-799 (RSS).

## References

1. Nelson PT, Dickson DW, Trojanowski JQ, Jack CR, Boyle PA, Arfanakis K, Rademakers R, Alafuzoff I, Attems J, Brayne C, Coyle-Gilchrist ITS, Chui HC, Fardo DW, Flanagan ME, Halliday G, Hokkanen SRK, Hunter S, Jicha GA, Katsumata Y, Kawas CH, Keene CD, Kovacs GG, Kukull WA, Levey AI, Makkinejad N, Montine TJ, Murayama S, Murray ME, Nag S, Rissman RA, Seeley WW, Sperling RA, White CL 3rd, Yu L, Schneider JA. Limbic-predominant age-related TDP-43 encephalopathy (LATE): consensus working group report. Brain. 2019 Jun 1;142(6):1503–1527.

2. Schneider JA. Neuropathology of Dementia Disorders. Continuum (Minneap Minn). 2022 Jun 1;28(3):834–851.

3. Ling JP, Pletnikova O, Troncoso JC, Wong PC. TDP-43 repression of nonconserved cryptic exons is compromised in ALS-FTD. Science 2015;349(6248):650–655.

4. Melamed Ze, López-Erauskin J, Baughn MW, Zhang O, Drenner K, Sun Y, Freyermuth F, McMahon MA, Beccari MS, Artates JW, Ohkubo T, Rodriguez M, Lin N, Wu D, Bennett CF, Rigo F, Da Cruz S and others. Premature polyadenylation-mediated loss of stathmin-2 is a hallmark of TDP-43-dependent neurodegeneration. Nature Neuroscience 2019;22(2):180–190.

5. Baughn MW, Melamed Z, López-Erauskin J, Beccari MS, Ling K, Zuberi A, Presa M, Gonzalo-Gil E, Maimon R, Vazquez-Sanchez S, Chaturvedi S, Bravo-Hernández M, Taupin V, Moore S, Artates JW, Acks E, Ndayambaje IS, Agra de Almeida Quadros AR, Jafar-Nejad P, Rigo F, Bennett CF, Lutz C, Lagier-Tourenne C, Cleveland DW. Mechanism of STMN2 cryptic splice-polyadenylation and its correction for TDP-43 proteinopathies. Science. 2023 Mar 17;379(6637):1140–1149.

6. Klim JR, Williams LA, Limone F, Guerra San Juan I, Davis-Dusenbery BN, Mordes DA, Burberry A, Steinbaugh MJ, Gamage KK, Kirchner R, Moccia R, Cassel SH, Chen K, Wainger BJ, Woolf CJ, Eggan K. ALS-implicated protein TDP-43 sustains levels of STMN2, a mediator of motor neuron growth and repair. Nature Neuroscience 2019;22(2):167–179.

7. Akiyama T, Koike Y, Petrucelli L, Gitler AD. Cracking the cryptic code in amyotrophic lateral sclerosis and frontotemporal dementia: Towards therapeutic targets and biomarkers. Clinical and Translational Medicine 2022;12(5):e818.

8. Prudencio M, Humphrey J, Pickles S, Brown A-L, Hill SE, Kachergus JM, Shi J, Heckman MG, Spiegel MR, Cook C, Song Y, Yue M, Daughrity LM, Carlomagno Y, Jansen-West K, de Castro CF, DeTure M and others. Truncated stathmin-2 is a marker of TDP-43 pathology in frontotemporal dementia. The Journal of Clinical Investigation 2020;130(11).

9. Vanden Broeck L, Naval-Sánchez M, Adachi Y, Diaper D, Dourlen P, Chapuis J, Kleinberger G, Gistelinck M, Van Broeckhoven C, Lambert JC, Hirth F, Aerts S, Callaerts P, Dermaut B. TDP-43 loss-of-function causes neuronal loss due to defective steroid receptor-mediated gene program switching in Drosophila. Cell Rep. 2013 Jan 31;3(1):160–72.

10. Diaper DC, Adachi Y, Sutcliffe B, Humphrey DM, Elliott CJ, Stepto A, Ludlow ZN, Vanden Broeck L, Callaerts P, Dermaut B, Al-Chalabi A, Shaw CE, Robinson IM, Hirth F. Loss and gain of Drosophila TDP-43 impair synaptic efficacy and motor control leading to age-related neurodegeneration by loss-of-function phenotypes. Hum Mol Genet. 2013 Apr 15;22(8):1539–57.

11. Wu LS, Cheng WC, Chen CY, Wu MC, Wang YC, Tseng YH, Chuang TJ, Shen CJ. Transcriptomopathies of pre- and post-symptomatic frontotemporal dementia-like mice with TDP-43 depletion in forebrain neurons. Acta Neuropathol Commun. 2019 Mar 29;7(1):50.

12. Pickles S, Gendron TF, Koike Y, Yue M, Song Y, Kachergus JM, Shi J, DeTure M, Thompson EA, Oskarsson B, Graff-Radford NR, Boeve BF, Petersen RC, Wszolek ZK, Josephs KA, Dickson DW, Petrucelli L, Cook CN, Prudencio M. Evidence of cerebellar TDP-43 loss-of-function in FTLD-TDP. Acta Neuropathol Commun. 2022 Jul 25;10(1):107.

13. Ebstein SY, Yagudayeva I, Shneider NA. Mutant TDP-43 Causes Early-Stage Dose-Dependent Motor Neuron Degeneration in a TARDBP Knockin Mouse Model of ALS. Cell Rep. 2019 Jan 8;26(2):364–373.e4.

14. Altman T, Ionescu A, Ibraheem A, Priesmann D, Gradus-Pery T, Farberov L, Alexandra G, Shelestovich N, Dafinca R, Shomron N, Rage F, Talbot K, Ward ME, Dori A, Krüger M, Perlson E. Axonal TDP-43 condensates drive neuromuscular junction disruption through inhibition of local synthesis of nuclear encoded mitochondrial proteins. Nat Commun. 2021 Nov 25;12(1):6914.

15. Yusuff T, Chang YC, Sang TK, Jackson GR, Chatterjee S. Codon-optimized TDP-43 mediates neurodegeneration in a Drosophila model of ALS/FTLD. Front Genet. 2023 Mar 9;14:881638.

16. Henstridge CM, Sideris DI, Carroll E, Rotariu S, Salomonsson S, Tzioras M, McKenzie CA, Smith C, von Arnim CAF, Ludolph AC, Lulé D, Leighton D, Warner J, Cleary E, Newton J, Swingler R, Chandran S, Gillingwater TH, Abrahams S, Spires-Jones TL. Synapse loss in the prefrontal cortex is associated with cognitive decline in amyotrophic lateral sclerosis. Acta Neuropathol. 2018 Feb;135(2):213–226.

17. Gregory JM, McDade K, Bak TH, Pal S, Chandran S, Smith C, Abrahams S. Executive, language and fluency dysfunction are markers of localised TDP-43 cerebral pathology in non-demented ALS. Journal of Neurology, Neurosurgery & Psychiatry 2020;91(2):149.

18. Banerjee P, Elliott E, Rifai OM, O’Shaughnessy J, McDade K, Abrahams S, Chandran S, Smith C, Gregory JM. NLRP3 inflammasome as a key molecular target underlying cognitive resilience in amyotrophic lateral sclerosis. J Pathol. 2022 Mar;256(3):262–268.

19. Gregory JM, Elliott E, McDade K, Bak T, Pal S, Chandran S, Abrahams S, Smith C. Neuronal clusterin expression is associated with cognitive protection in amyotrophic lateral sclerosis. Neuropathol Appl Neurobiol. 2020 Apr;46(3):255–263.

20. Zacco E, Kantelberg O, Milanetti E, Armaos A, Panei FP, Gregory J, Jeacock K, Clarke DJ, Chandran S, Ruocco G, Gustincich S, Horrocks MH, Pastore A, Tartaglia GG. Probing TDP-43 condensation using an in silico designed aptamer. Nat Commun. 2022 Jun 23;13(1):3306.

21. Bankhead P, Loughrey MB, Fernández JA, Dombrowski Y, McArt DG, Dunne PD, McQuaid S, Gray RT, Murray LJ, Coleman HG, James JA, Salto-Tellez M, Hamilton PW. QuPath: Open source software for digital pathology image analysis. Sci Rep. 2017 Dec 4;7(1):16878.

22. Wickham, H. ggplot2: Elegant Graphics for Data Analysis. Springer-Verlag New York 2016.

23. Chou CC, Zhang Y, Umoh ME, Vaughan SW, Lorenzini I, Liu F, Sayegh M, Donlin-Asp PG, Chen YH, Duong DM, Seyfried NT, Powers MA, Kukar T, Hales CM, Gearing M, Cairns NJ, Boylan KB, Dickson DW, Rademakers R, Zhang YJ, Petrucelli L, Sattler R, Zarnescu DC, Glass JD, Rossoll W. TDP-43 pathology disrupts nuclear pore complexes and nucleocytoplasmic transport in ALS/FTD. Nat Neurosci. 2018 Feb;21(2):228–239.

24. Yabata, H.; Riku, Y.; Miyahara, H.; Akagi, A.; Sone, J.; Urushitani, M.; Yoshida, M.; Iwasaki, Y. Nuclear Expression of TDP-43 Is Linked with Morphology and Ubiquitylation of Cytoplasmic Aggregates in Amyotrophic Lateral Sclerosis. Int. J. Mol. Sci. 2023, 24, 12176.

25. Coyne AN, Baskerville V, Zaepfel BL, Dickson DW, Rigo F, Bennett F, Lusk CP, Rothstein JD. Nuclear accumulation of CHMP7 initiates nuclear pore complex injury and subsequent TDP-43 dysfunction in sporadic and familial ALS. Sci Transl Med. 2021 Jul 28;13(604):eabe1923.

26. Zhou J, Rossi J. Aptamers as targeted therapeutics: current potential and challenges. Nat Rev Drug Discov. 2017 Mar;16(3):181–202. doi: 10.1038/nrd.2016.199. Epub 2016 Nov 3. Erratum in: Nat Rev Drug Discov. 2017 Jun;16(6):440.

27. Kim ST, Kim HG, Kim YM, Han HS, Cho JH, Lim SC, Lee T, Jahng GH. An aptamer-based magnetic resonance imaging contrast agent for detecting oligomeric amyloid-β in the brain of an Alzheimer’s disease mouse model. NMR Biomed. 2023 Mar;36(3):e4862.

28. Liu D, Xia Q, Ding D, Tan W. Radiolabeling of functional oligonucleotides for molecular imaging. Front Bioeng Biotechnol. 2022 Aug 19;10:986412.

