## Supplemental figures for "RNA aptamer reveals nuclear TDP-43 pathology is an early aggregation event that coincides with *STMN-2* cryptic splicing and precedes clinical manifestation in ALS"

**Supplementary Information**

**Supplementary Figures:**

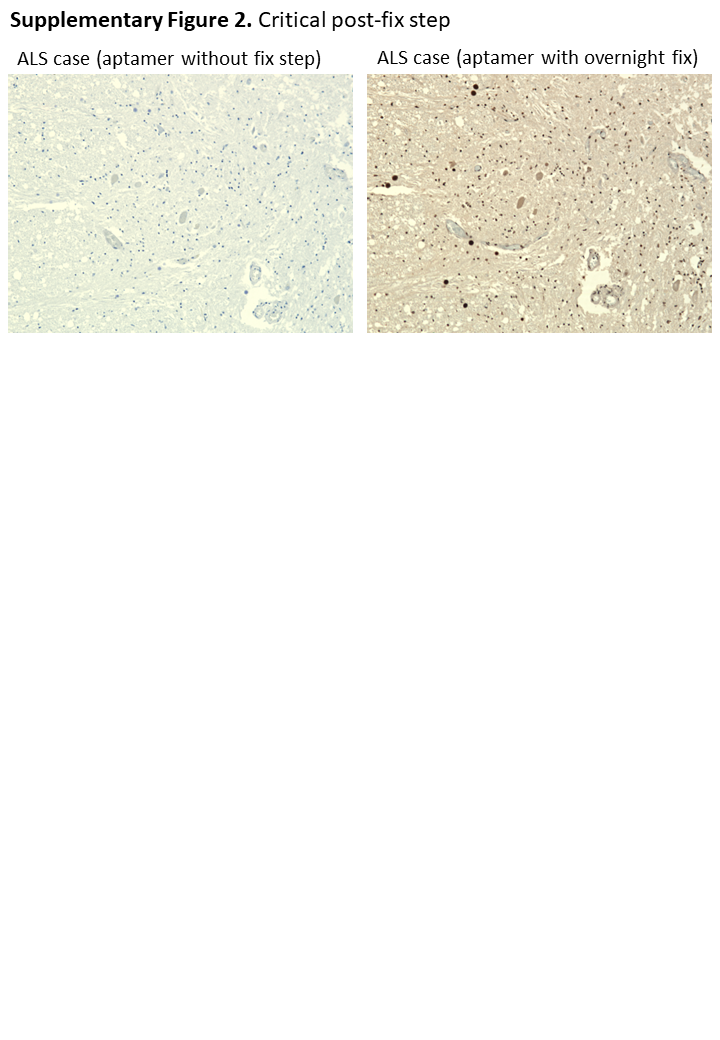
**Supplementary Figure 1. Critical Post-Fix Step**

Serial sections from the spinal cord of a patient with sporadic ALS stained without (left image) and with (right image) the critical post-fix step (incubation with 4% PFA overnight after aptamer incubation) demonstrate that it is necessary to immobilise the aptamer on to the target using formalin cross links to enable immunodetection. Scale bar is 200 µM.

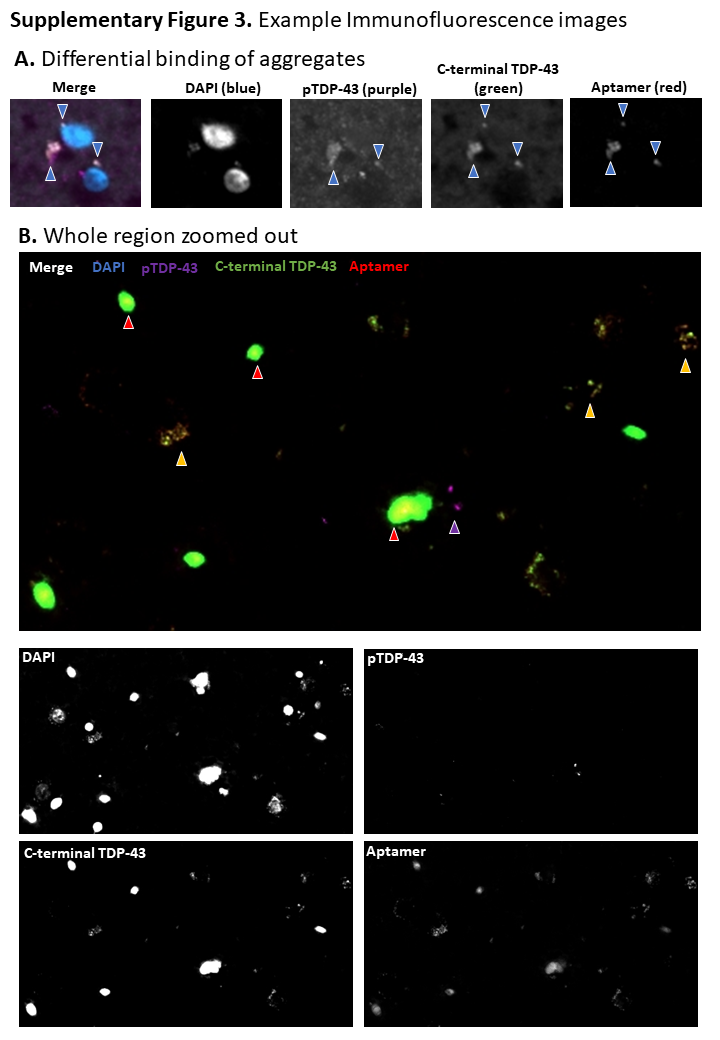

**Supplementary Figure 2.**

Immunofluorescent images showing A. Two neuronal cells containing cytoplasmic TDP-43 aggregates that stain for all three antibody markers (pTDP-43 in purple, c-terminal TDP-43 in green and TDP-43^APT^ in red). Blue arrows indicate an aggregate present in all three channels. Scale bar is 5 µm. **B**. Zoomed out region with extensive TDP-43 aggregation - immunofluorescent image demonstrating the variation in aggregation events, with evidence of pTDP-43 immunoreactive aggregates (purple arrowhead) alone as well as co-incident immunoreactivity in some aggregates for c-terminal antibody and TDP-43^APT^ (yellow arrowhead). There are also multiple examples of TDP-43^APT^ staining within the nucleus which is obscured by normal c-terminal TDP-43 antibody staining. Scale bar is 10 µm.

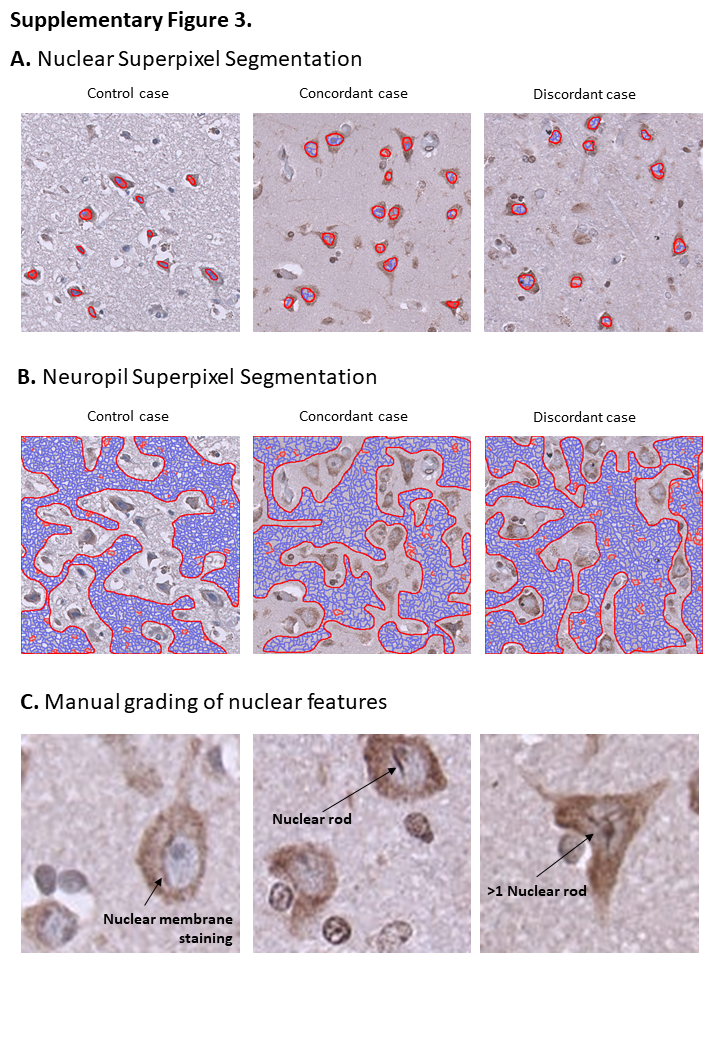

**Supplementary Figure 3.**

Example images of nuclear superpixel segmentation (A) and neuropil superpixel segmentation (B) performed using QuPath software and example images of nuclear features graded by blinded manual assessment (C).
